# Differentiation-Dependent Proximity Proteomics Identifies Novel Host Factors Linked to HPV16 E2 Function

**DOI:** 10.1101/2025.10.21.683683

**Authors:** Claire D. James, Aya Youssef, Apurva T. Prabhakar, Jenny D Roe, Elinor Lu, Austin Witt, S Giri, Molly L. Bristol, Phoebe Bridy, Xu Wang, Arjun Rijal, Charles Lyons, Iain M. Morgan

## Abstract

Human papillomavirus 16 (HPV16) is a causative agent of oropharyngeal, cervical and anogenital cancers. The viral E2 protein is essential for viral genome replication, transcriptional regulation, episome maintenance, and activation of the host DNA damage response. Despite its central role, the full network of HPV16 E2 interactions with host proteins remains incompletely defined, particularly under differentiating conditions which support the complete viral life cycle. In this study, we used TurboID-based proximity labeling to characterize the interactome of HPV16 E2 and known host partner protein TOPBP1, in both undifferentiated monolayer and differentiating keratinocytes. We generated stable keratinocyte lines expressing doxycycline-inducible TurboID-tagged HPV16 E2 and confirmed that the tagged protein retained transcriptional, replicative, and DNA damage-inducing functions. Mass spectrometry analysis of streptavidin-enriched proteins identified both known and novel E2-associated host factors, including chromatin regulators, DNA repair proteins, and nucleolar components. Comparative analysis revealed a substantial overlap between E2 and TOPBP1 interactomes, and *in situ* validation by proximity ligation assay identified nucleolin (NCL) as a differentiation-dependent factor whose interaction with E2 is stabilized by TOPBP1. Functional studies demonstrated that NCL is required for episomal genome maintenance, highlighting a cooperative E2–TOPBP1– NCL axis critical for viral genome stability during differentiation. These findings provide a comprehensive view of the E2-associated protein landscape in stratified epithelial cells and reveal a mechanistic pathway through which HPV16 co-opts host factors to support genome maintenance, productive replication, and persistence.

**Importance:** Human papillomaviruses (HPVs) establish persistent infections in stratified epithelia and rely on host DNA damage and repair factors to support their replication. The E2 protein is central to viral genome replication and maintenance, and depends heavily on its interaction with the host factor TOPBP1 for these functions. Here, we define the E2 and TOPBP1 interactomes in differentiating keratinocytes, and identify nucleolin (NCL) a critical differentiation- and TOPBP1-dependent E2 partner required for episomal genome stability. These findings expand the understanding of how HPV16 coordinates viral replication with host chromatin and DNA repair networks, uncovering a cooperative E2–TOPBP1–NCL axis that may represent a new target for antiviral intervention.

## Introduction

Human Papillomavirus type 16 (HPV16) is the predominant oncogenic HPV type associated with anogenital and oropharyngeal cancers, the latter of which is rising in incidence in the United states (1–3). Globally, HPV16 accounts for the majority of HPV-positive head and neck squamous cell carcinomas (HNSCC), representing a major clinical and public health concern (4). Unlike cervical cancers, in which viral genomes are integrated in 90% of cases, a substantial fraction (30-60%) of HPV16-positive oropharyngeal cancers retain episomal genomes (5–7). Viral Integration disrupts the E2 open reading frame, abrogating its regulatory function and leading to unchecked E6/E7 activity (7, 8). In contrast, the retention of episomal genomes preserves E2, suggesting that E2 may actively shape tumor biology in a subset of HPV16-driven oropharyngeal cancers (7, 8). E2 is a multifunctional viral regulator which controls viral transcription, mediates initiation of viral DNA replication, tethers genomes to mitotic chromosomes, and engages the host DNA damage response (DDR), a pathway that facilitates productive viral replication in differentiating epithelial cells (9–11). The HPV life cycle is tightly linked to the differentiation program of stratified epithelium (12). Viral genomes are maintained at low copy numbers in the dividing basal cells of the epithelium. As these infected cells undergo terminal differentiation, viral genome amplification is triggered, along with late gene expression and virion production (12); these processes rely on the ability of E2 to engage with cellular proteins. Several E2-binding proteins have been described to date, including BRD4, p300, ChlR1, and TOPBP1, and these studies have provided important mechanistic insights into viral replication and gene regulation (13–22). Much of this work has been conducted in transformed or non-epithelial systems, leaving significant gaps in our understanding of the E2-host interactome, particularly under physiologically relevant conditions. Importantly, prior analyses have not accounted for the differentiation-dependent nature of the HPV life cycle, leaving unanswered how E2 rewires its interactions across epithelial states. For the first time, we demonstrate that E2 engages with both common and differentiation-specific host proteins, reflecting its versatile role in coordinating viral processes as cells transition from basal proliferation to terminal differentiation.

In this study, we developed a doxycycline-inducible proximity labeling model in human keratinocytes using TurboID-tagged HPV16 E2. TurboID enables *in situ* biotinylation of proteins in the immediate vicinity of the bait, allowing unbiased identification of both stable and transient interactors (23). By inducing TurboID-E2 under both monolayer and differentiation conditions, we captured the interactome of E2 across epithelial states that mimic the physiological environment of the viral life cycle. In parallel, we applied the same approach to TOPBP1, an established E2 interactor and multifunctional scaffold protein involved in DNA replication and repair (19–22, 24–28), to enable direct comparison of their interactomes. Through mass spectrometry and functional validation, we characterize the host protein partners of E2 and TOPBP1, uncovering both shared and unique interaction networks that may contribute to HPV16 genome maintenance, productive replication, long-term persistence and oncogenesis. Our analyses revealed a spectrum of host protein partners, including factors involved in chromatin regulation, DNA replication, and the DNA damage response. Comparative mapping of the E2 and TOPBP1 networks uncovered both shared and unique interactors, suggesting distinct as well as cooperative roles for these proteins during viral genome maintenance and amplification. Notably, we identify an interaction between E2 and nucleolin (NCL), a host protein known as a histone chaperone, but with multiple cellular roles including chromatin remodeling, RNA metabolism, and DNA repair, replication, and recombination (29). This interaction was most prominent under differentiating conditions, suggesting a role during the amplification phase of the viral life cycle. Additionally, E2 binding to TOPBP1 enhanced the TOPBP1–NCL interaction, and functional studies demonstrated that NCL is required for episomal genome maintenance, identifying it as a novel host factor supporting HPV16 persistence. Our findings reveal that the E2 interactome is not static, but is dependent on epithelial context, providing new insight into how HPV16 exploits host pathways to balance genome maintenance, productive replication, and persistence.

## Results

### Generation of inducible TurboID-tagged HPV16 E2 and TOPBP1 keratinocyte models

To investigate the HPV16 E2 interactome in a physiologically relevant epithelial background, we employed the biotin ligase, TurboID (23, 30), to label and enrich proteins in close proximity to the viral E2 protein. The E2 open reading frame was cloned into a doxycycline-inducible TurboID expression construct (pCW57.1_V5_TurboID_FLAG_Nt), resulting in a fusion protein with an N-terminal TurboID-3×FLAG tag and a C-terminal V5 tag (Figure 1A). The same strategy was used to generate a TurboID-tagged version of the known E2 interactor, TOPBP1, to enable comparison between the two protein interactomes. Figure 1B describes the strategy for the identification of E2 and TOPBP1 interacting partners: Lentiviral transduction followed by blasticidin selection were used to generate stable N/Tert-1 and N/Tert-1+HPV16 (containing the full HPV16 genome) keratinocyte lines carrying TurboID-E2 or TurboID-TOPBP1 constructs. We have described N/Tert-1+HPV16 previously; it retains many aspects of the viral life cycle (8, 31). Doxycycline-induction of the TurboID-tagged proteins was confirmed by western blotting in both monolayer and differentiating (Ca²⁺-treated) cells. Both E2 and TOPBP1 fusion proteins were strongly expressed following treatment with doxycycline for 48 hours. Figure 1C shows N/Tert-1 (top 3 panels) and N/Tert-1HPV16 (lower 3 panels) lines stably expressing the inducible TurboID-tagged E2 vector. Tagged E2 expression induced by doxycycline was confirmed in both monolayer and calcium-treated cells, utilizing antibodies to E2 and to the TurboID tag (BirA) (Figure 1C, induced lanes 2 and 4 compared to uninduced lanes 1 and 3). In N/Tert-1+HPV16, endogenous E2 is visible as the lower band (E2 endo), and exogenous TurboID-E2 as the higher band (E2 Exo); the TurboID tag adds 35kDa (32). Biotinylation and successful streptavidin pulldown of TurboID-tagged E2 was confirmed in monolayer and calcium-treated N/Tert-1 and N/Tert-1+HPV16 (Figure 1C, doxycycline induced lanes 6 and 8 compared to uninduced lanes 5 and 7).

**Figure 1:**
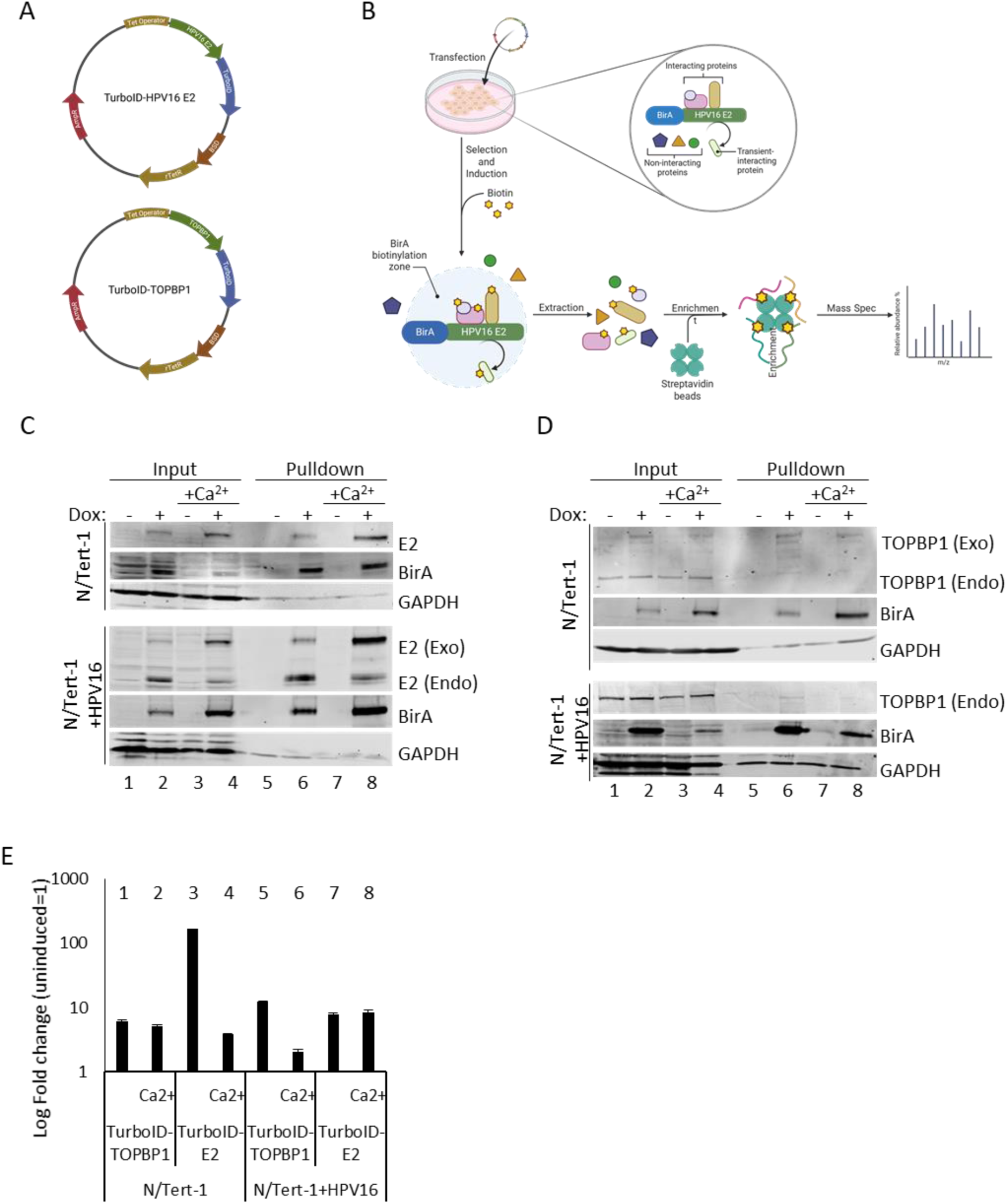
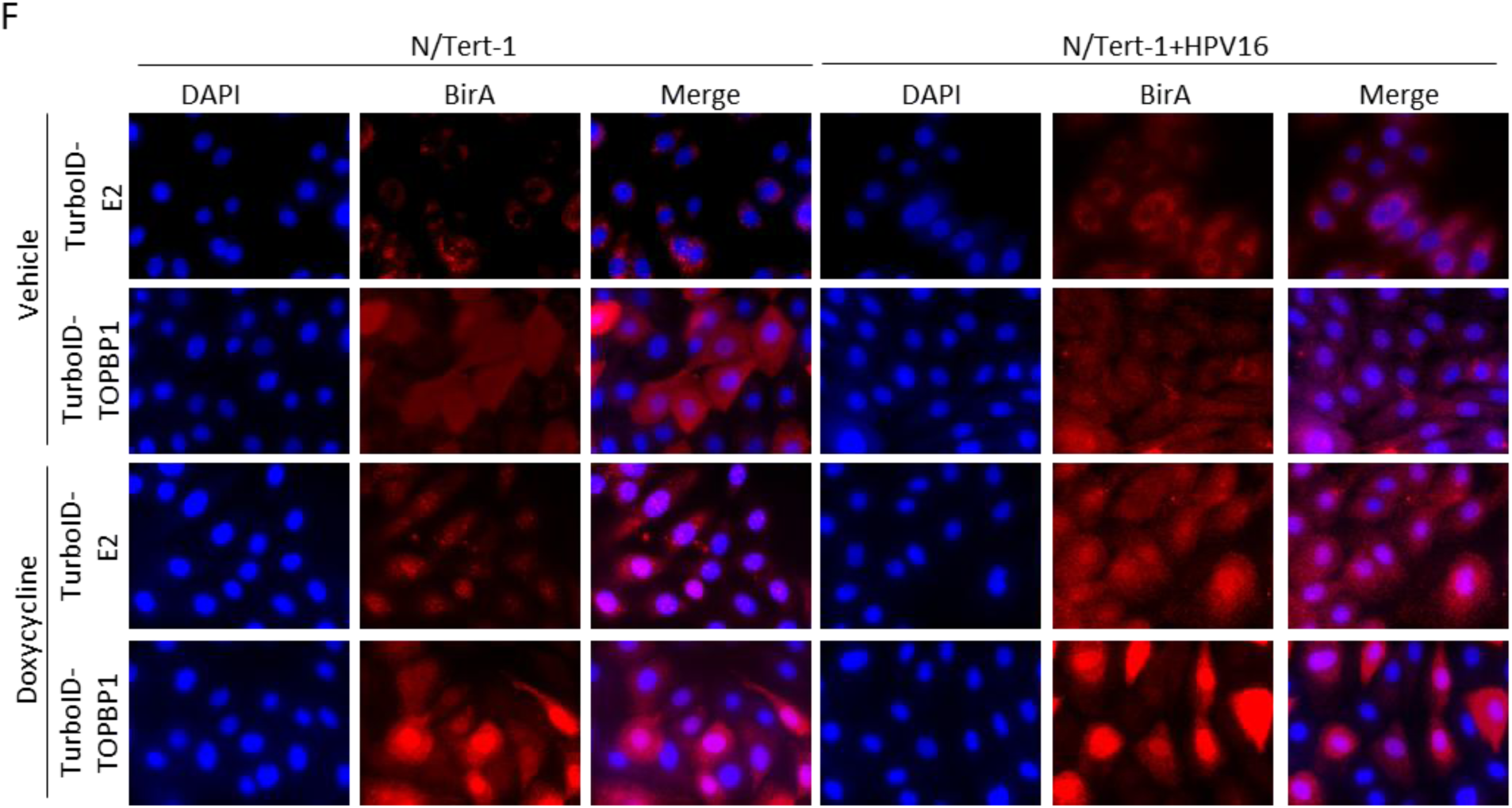
Generation of stable N/Tert-1 and N/Tert-1+HPV16 lines expressing inducible doxycycline-inducible Turbo-ID tagged proteins. A. Schematic maps of doxycycline-inducible constructs encoding TurboID-tagged HPV16 E2 and TOPBP1. Created with Biorender B. Overview of the experimental workflow. Constructs were introduced into N/Tert-1 and N/Tert-1+HPV16 cells, followed by blasticidin selection to generate stable lines. Upon doxycycline induction, the promiscuous biotin ligase TurboID biotinylates proteins in proximity to the tagged bait protein. Biotinylated proteins were enriched using streptavidin magnetic beads and identified by mass spectrometry. C. Western blot analysis confirmed inducible expression of TurboID-tagged E2 in N/Tert-1 and N/Tert-1+HPV16 cells under both monolayer and differentiating (1.5 mM CaCl₂) conditions. Streptavidin-bead pulldown confirmed successful enrichment of the bait protein. D. Western blot analysis confirmed inducible expression of TurboID-tagged TOPBP1 in N/Tert-1 and N/Tert-1+HPV16 cells under both monolayer and differentiating (1.5 mM CaCl₂) conditions. Streptavidin-bead pulldown confirmed successful enrichment of the bait proteins. E. qRT-PCR using primers targeting the BirA confirmed induction of TurboID expression following doxycycline treatment. Fold changes were calculated relative to GAPDH and normalized to uninduced controls. Error bars represent standard deviation from biological replicates. F. Immunofluorescence of N/Tert-1 and N/Tert-1+HPV16 grown on glass coverslips confirmed nuclear localization of TurboID-tagged proteins using a BirA-specific antibody. Images captured at 100× magnification.

Similarly, tagged TOPBP1 was induced in N/Tert-1 and N/Tert-1+HPV16 grown in both monolayer and calcium, and expression confirmed by western blot with BirA and TOPBP1 antibodies (Figure 1D, induced lanes 2 and 4 compared to uninduced lanes 1 and 3). Pulldown of biotinylated TOPBP1 with streptavidin beads was confirmed in both growing and differentiating conditions (Figure 1D, doxycycline-induced lanes 6 and 8 compared to uninduced lanes 5 and 7).

To further validate tagged protein induction, qRT-PCR targeting the TurboID (BirA) sequence was performed on the N/Tert-1 and N/Tert-1+HPV16 grown in monolayer and under differentiating conditions (Ca^2+^). Across all conditions, doxycycline-treated cells exhibited a significant increase in TurboID transcript levels compared to uninduced controls, normalized to GAPDH (Figure 1E). In N/Tert-1 Cells, turboID-tagged TOPBP1 transcripts increased approximately 5-fold under monolayer conditions and 6-fold under calcium induced differentiation (Figure 1E lanes 1 and 2). In N/Tert-1+HPV16, a 12-fold induction was observed in monolayer conditions and a 2-fold induction under differentiating conditions (lanes 5 and 6). Doxycycline treatment induced TurboID-tagged E2 more robustly, with over a 100-fold increase in N/Tert-1 monolayers and a 4-fold increase under differentiation (lanes 3 and 4). In N/Tert-1+HPV16 cells, E2 transcripts increased approximately 8-fold in both conditions (lanes 7 and 8). These transcript-level increases, together with the corresponding protein induction observed by western blot, confirm that the doxycycline-inducible TurboID tagging system effectively drives expression of both TOPBP1 and E2 fusion proteins across the different epithelial states.

Immunofluorescence staining with an anti-BirA antibody confirmed nuclear localization of TurboID-tagged proteins in both N/Tert-1 and N/Tert-1+HPV16 cells (Figure 1F), consistent with the known nuclear localization of E2 and TOPBP1(21). There is also clear staining in the cytoplasm following induction but there is a concentration in the nuclei. Together, these results establish a robust, inducible platform for mapping E2- and TOPBP1-proximal proteomes in keratinocyte models that can recapitulate key aspects of HPV biology.

#### TurboID-tagged proteins retain function

To determine whether TurboID-tagged proteins retained their expected biological activities, we validated E2 and TOPBP1 function in keratinocytes (Figure 2). We first assessed the activity of TurboID-tagged E2 in transcriptional and replication assays. In luciferase-based reporter assays performed in N/Tert-1 cells, TurboID-E2 activated a reporter with 6 E2 DNA binding sites located upstream from the HSV-1 Thymindine Kinase promoter (TK) (pTK6E2), consistent with known E2 functions in transcriptional activation and repression (33–37) (Figure 2 A). TurboID-TOPBP1 was included as a negative control and showed no effect on either activation or repression (lanes 1 and 3). Upon doxycycline induction, E2 expression resulted in robust activation of the pTK6E2 reporter (lane 4 vs. lane 2), confirming that the TurboID-tagged E2 retains its transcription related functions.

**Figure 2:**
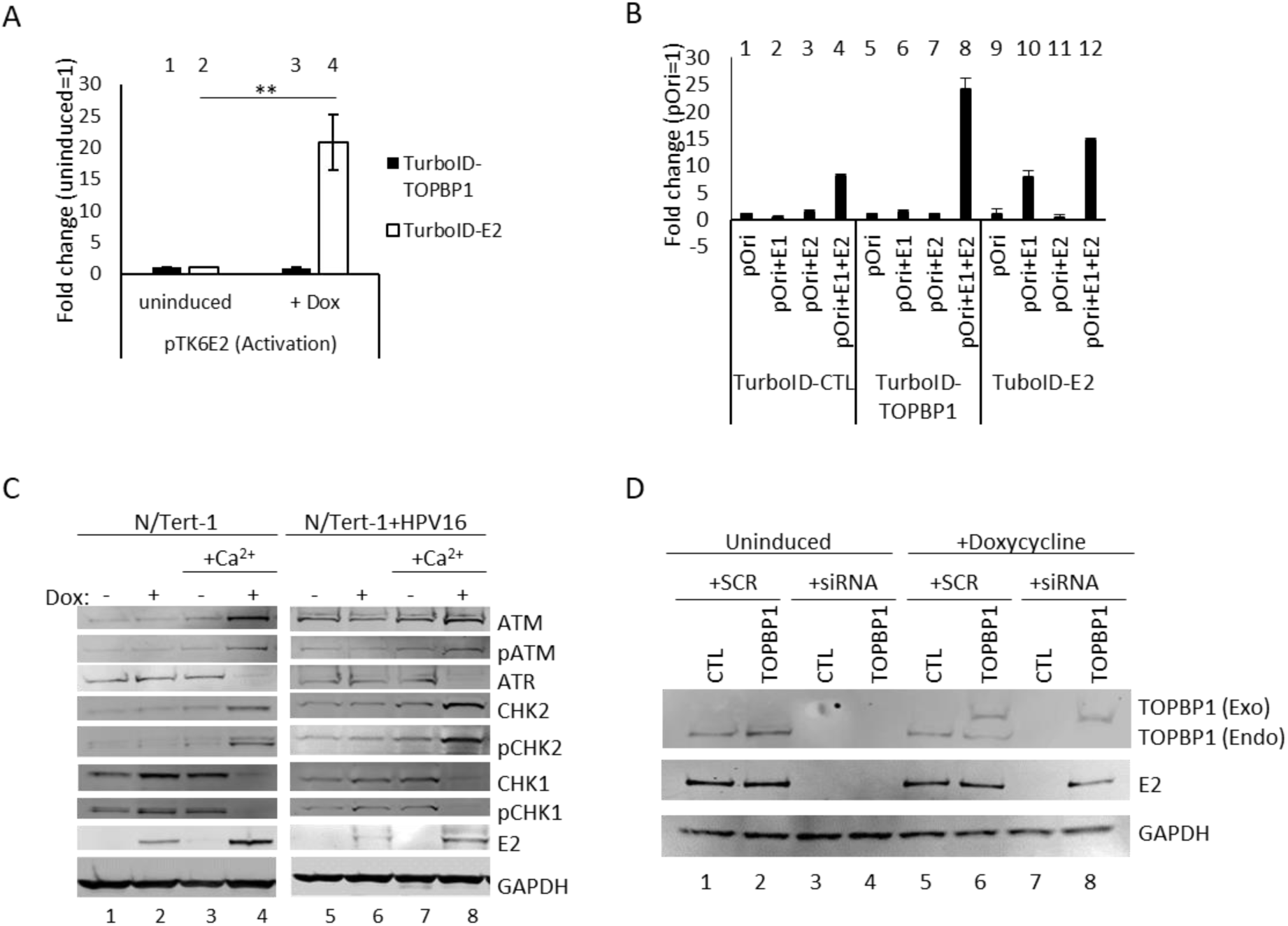
TurboID-tagged E2 and TOPBP1 retain functional activity. A. Luciferase-based transcriptional assays in N/Tert-1 cells demonstrate that doxycycline-induced TurboID-E2 retains the ability to activate transcription. Bars represent the mean of three biological replicates; error bars indicate standard error of the mean (SEM). Statistical significance was assessed using unpaired t-tests; **p < 0.01. B. DNA replication assays in a C33a background confirm that TurboID-tagged E2 supports viral origin-dependent replication upon induction. Bars represent the mean of three biological replicates; error bars indicate standard error of the mean (SEM). C. Western blot analysis of differentiating N/Tert-1 and N/Tert-1+HPV16 cells shows activation of the DNA damage response (DDR) following TurboID-E2 expression. Upon doxycycline induction, increased levels of total ATM and phosphorylated CHK2 were observed, accompanied by decreased levels of ATR and CHK1, under differentiating conditions. D. N/Tert-1+HPV16 cells stably expressing either TurboID vector control (CTL) or TurboID-TOPBP1 (TOPBP1) were transfected with scrambled (SCR) or TOPBP1-targeting siRNA (+siRNA) and cultured under uninduced conditions or doxycycline-induced (+Dox) conditions. siRNA targeted the 3′ UTR to silence endogenous TOPBP1. Western blot analysis shows endogenous TOPBP1 (lower band) and exogenous TurboID-tagged TOPBP1 (upper band). Silencing TOPBP1 reduced E2 protein levels, while induction of TurboID-TOPBP1 rescued E2 stability, confirming that the tagged protein retains functional activity.

To assess the ability of TurboID-E2 to support origin-dependent DNA replication, C33A cells stably expressing TurboID-tagged E2 and TOPBP1 (Supplementary Figure 1) were co-transfected with E1 and a plasmid containing the HPV16 origin, and treated with doxycycline to induce expression. Cells containing the empty vector and TurboID-tagged TOPBP1 (lanes 1-8) showed no background replication when transfected with pOri, E1 and E2 individually (CTL lanes 1, 2 and 3; TOPBP1 lanes 5, 6 and 7), and successful replication when transfected with the complete replication complex (lanes 4 and 8), indicating that doxycycline treatment did not impede viral replication (Figure 2C). Induction of TurboID-E2 expression significantly increased replication levels in cells transfected with pOri and E1 (lane 10), whereas transfecting with pOri ond E2 alone (lanes 9 and 11) showed no induction of replication, altogether demonstrating that the E2 fusion protein is competent for replication (Figure 2C).

Given the known role of E2 in activating the host DNA damage response (DDR) during differentiation (11), we examined DDR signaling in cells growing in monolayer and in differentiating conditions (Figure 2D). Upon TurboID-E2 induction and calcium-induced differentiation, western blot analysis of N/Tert-1 (lanes 1-4) and N/Tert-1+HPV16 (lanes 5-8) revealed elevated ATM and CHK2 activation, as evidenced by increased levels of their phosphorylated forms, and a concurrent reduction in ATR and CHK1 pathway activity (Figure 2D, compare lanes 4 and 8 to 3 and 7). This pattern suggests that E2 expression promotes a shift in DDR signaling towards an ATM/CHK2-dominated response in differentiating keratinocytes, consistent with its role in supporting viral genome amplification (37). This same pattern exists in multiple models of the virus lifecycle and HPV-driven carcinogenesis (11).

To confirm TurboID-TOPBP1 function, the protein was induced in N/Tert-1+HPV16 cells and endogenous TOPBP1 subjected to siRNA-mediated knockdown targeting the 3′ UTR (Figure 2E). Cells expressing the empty turboID vector (CTL) were treated in parallel with TurboID-TOPBP1 expressing cells. Silencing of endogenous TOPBP1 resulted in a marked reduction in E2 protein levels (lanes 3,4 and 7), consistent with previous reports that TOPBP1 stabilizes E2 and supports its activity during the viral life cycle (20). When TurboID-TOPBP1 was induced under conditions of endogenous TOPBP1 depletion (lane 8), E2 stability was maintained, demonstrating that the exogenous TOPBP1 retains functional activity.

Together, these results confirm that TurboID-tagged E2 and TOPBP1 proteins retain key functional properties, including transcriptional regulation, replication activity, and activation of the DDR, further supporting their use in proximity-dependent interactome profiling.

#### Proteomic analysis identified previously described E2 and TOPBP1 interactions

Having established expression and functionality of TurboID tagged E2 in TOPBP1 in keratinocytes, we moved to compare the interaction networks of HPV16 E2 and known cellular binding partner TOPBP1. We performed TurboID proximity labeling and mass spectrometry in stable N/Tert-1 keratinocyte lines expressing TurboID-tagged E2 or TOPBP1. Each bait was induced by doxycycline, and labeling was performed under both monolayer and differentiating (Ca²⁺-treated) conditions to model basal and suprabasal epithelial states, respectively. This approach allowed us to capture the interactomes of each protein in the context of epithelial stratification, an essential feature of the HPV16 life cycle. Streptavidin-enriched lysates from TurboID-E2–expressing cells demonstrated a diverse set of host proteins across both monolayer and differentiating conditions. Supplementary Table 1 provides quantitative mass spectrometry data for all identified protein interactors (normalized abundance). Supplementary Table 2 lists the condition-specific protein interactors for TurboID-E2 and TurboID-TOPBP1, including those unique to monolayer or differentiation. These datasets together provide a comprehensive resource for the HPV16 E2 and TOPBP1 proximity interactomes.

Proteomic analyses revealed that differentiation dramatically reduced the number of interactors for E2, from 2170 to 406 (an 81% reduction, Figure 3A). Similarly, TOPBP1 goes from 2176 interactors in monolayer to 372 under differentiating conditions (an 82% reduction, Figure 3B). For both E2 and TOPBP1, this demonstrates a shift in the protein environment in response to epithelial differentiation. When compared to a previously published E2 interactome (39) (Figure 3C), our dataset showed validation of several previously identified E2 interactors including BRD4, TOP1 and HNRNPA1 in the monolayer sample; these interactors were lost following differentiation (Supplementary Table 2)(13, 14, 39–41). Similarly, we identified proteins interacting with TOPBP1 that have been identified in previous proteomic analyses of TOPBP1, including MRE11, MDC1, TP53BP1 (24, 42–44) (Figure 3D). These results validate the specificity and functionality of the TurboID-based approach.

**Figure 3.**
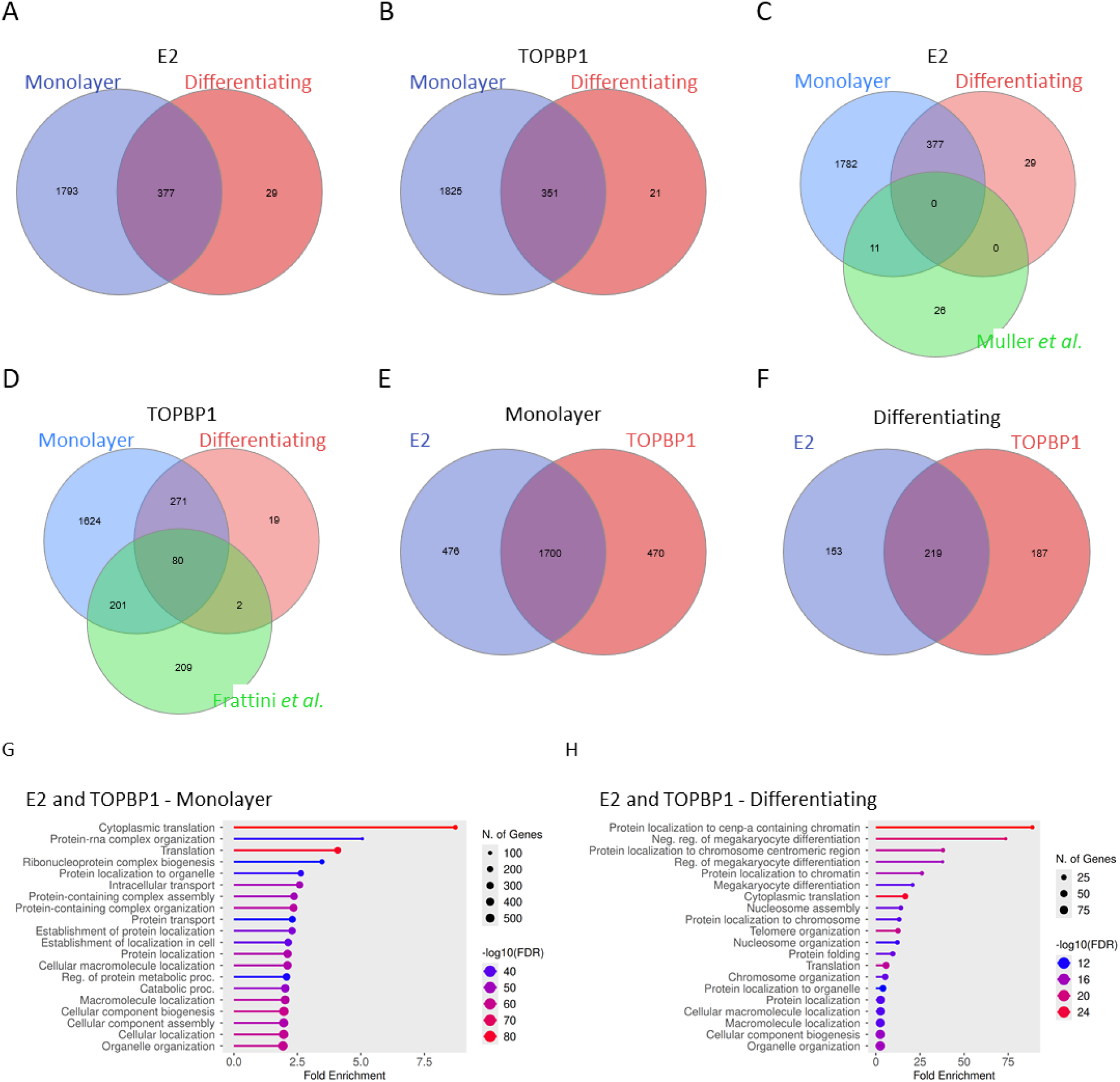
Proteomic analysis shows overlaps between E2 and TOPBP1 interactomes across growth conditions and with previously published datasets. A. Overlap of HPV16 E2 interactors identified under monolayer and differentiating conditions. B. Overlap of HPV16 E2 interactors identified under monolayer and differentiating conditions, in comparison with a previously published E2 interactome dataset from Muller *et al*. (40). C. Overlap of TOPBP1 interactors under monolayer and differentiating conditions. D. Overlap of TOPBP1 interactors under monolayer and differentiating conditions, in comparison with a previously published TOPBP1 interactome dataset from Frattini *et al*. (24). E. Comparison of E2 and TOPBP1 interactomes in monolayer conditions reveals substantial shared interactors. F. Comparison of E2 and TOPBP1 interactomes in differentiating conditions reveals substantial shared interactors. G. Top 20 enriched biological processes associated with HPV16 E2 and TOPBP1 shared interactors under monolayer conditions. H. Top 20 enriched biological processes associated with HPV16 E2 and TOPBP1 shared interactors under differentiating conditions. Venn diagrams were generated using InteractiVenn.net. Ontology analysis was performed using ShinyGO v0.829, with the *Homo sapiens* genome as background, applying an FDR cutoff of 0.05 and a minimum pathway size of 2. Circle size indicates the number of genes contributing to each term; color reflects statistical significance as –log₁₀(FDR).

Proteomic analyses revealed a substantial overlap between the E2 and TOPBP1 interactomes (Figure 3E and F) with 78% of E2 interactors identified in common with TOPBP1 in monolayer and 54% under differentiating conditions. The study identified a shared core set of interactors across conditions, suggesting context-dependent recruitment or stabilization of E2-TOPBP1 protein complexes during this process; both proteins are increased following differentiation (Figure 5E) (11). GO enrichment analysis of shared interactors between TOPBP1 and E2 in monolayer cultures revealed that the most significantly enriched pathway was cytoplasmic translation (FDR = 6.4 × 10⁻⁸⁶) (Figure 3G). Among the top 20 enriched pathways, the majority were associated with protein localization (FDR = 3.6 × 10⁻⁵⁴) and other translation-related processes. In contrast, under calcium-induced differentiation, the most significantly enriched GO category was protein localization to CENP-A–containing chromatin (FDR = 3.5 × 10⁻²⁷), with additional top pathways including nucleosome assembly (FDR = 3.2 × 10⁻¹⁴) and protein localization to the chromosome (FDR = 9.4 × 10⁻¹⁴) (Figure 3H). Together, these findings highlight the breadth of host processes engaged by E2 and TOPBP1 and indicate that their shared interactors are directed toward distinct biological pathways depending on the epithelial state. These results support a model in which E2 co-opts pre-existing host protein complexes through its interaction with TOPBP1, particularly within differentiated epithelial layers where viral genome amplification occurs.

#### HPV16 alters the TOPBP1 interactome in differentiating keratinocytes

To investigate how HPV16 influences the TOPBP1 interactome, we compared TurboID-TOPBP1–labeled proteins in N/Tert-1 cells with and without stably maintained HPV16 episomes (8, 31). In monolayer culture, the majority of TOPBP1-interacting proteins were shared between HPV16-negative and HPV16-positive cells. A small subset (9%) of proteins appeared to be selectively enriched in the HPV16-positive background (Figure 4A); these were associated with RNA polymerase and ribosome biogenesis (Figure 4B). GO enrichment analysis yielded a false discovery rate (FDR) of 0.02 and 0.04 for these categories, respectively. However, under differentiating conditions, the profile of TOPBP1 interactions shifted markedly. A larger fraction (54%) of proteins were enriched specifically in HPV16-positive cells (Figure 4C), indicating that viral protein expression (potentially E2) alters the associations of TOPBP1 in a differentiation-dependent manner. GO analysis of these HPV16-specific interactions under differentiating conditions revealed significant enrichment for translation-related processes, including proteosome (FDR=1.45×10^-10^), ribosome (FDR=1.45×10^-10^) and spliceosome (FDR=1.38×10^-8^) pathways (Figure 4D). These data suggest that HPV16 modulates TOPBP1-associated complexes in differentiating epithelial cells, potentially redirecting host protein synthesis, turnover and RNA processing machinery to support the productive phase of the viral life cycle.

**Figure 4.**
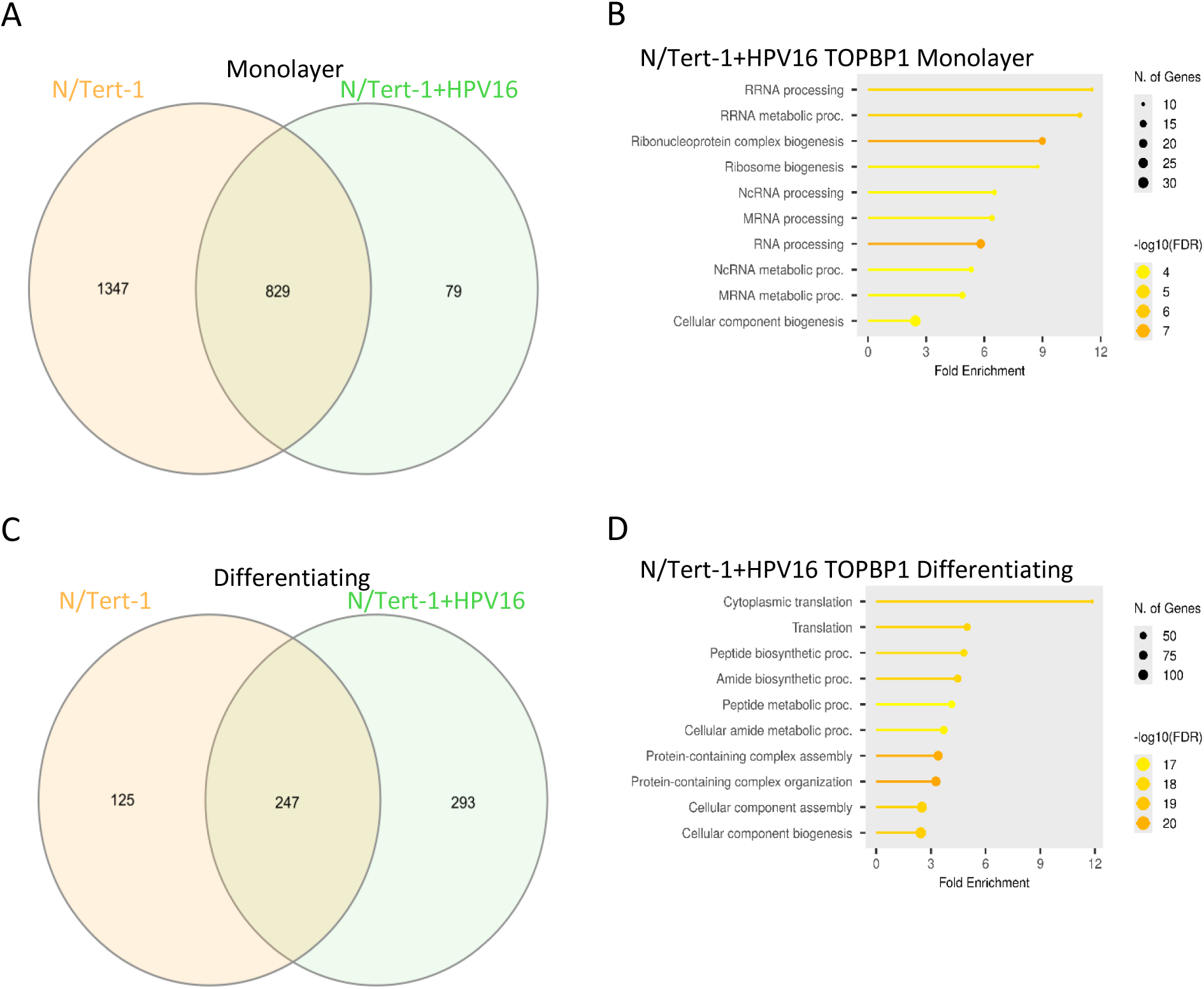
HPV16 alters the TOPBP1 interactome in differentiating keratinocyte. A. Venn diagram comparing TOPBP1-interacting proteins identified by TurboID proximity labeling in N/Tert-1 and N/Tert-1+HPV16 grown in non-differentiating conditions. B. Gene ontology (GO) enrichment analysis of HPV16-specific TOPBP1 interactors in monolayer-grown keratinocytes. C. Venn diagram comparing TOPBP1-interacting proteins identified by TurboID proximity labeling in N/Tert-1 and N/Tert-1+HPV16 keratinocytes cultured under differentiating conditions. D. Gene ontology (GO) enrichment analysis of HPV16-specific TOPBP1 interactors in differentiating cells. Venn diagrams were generated using InteractiVenn.net. Ontology analysis was performed using ShinyGO v0.829, with the *Homo sapiens* genome as background, applying an FDR cutoff of 0.05 and a minimum pathway size of 2. Circle size indicates the number of genes contributing to each term; color reflects statistical significance as –log₁₀(FDR).

#### E2 interacts with nucleolin in a differentiation- and TOPBP1-dependent manner

Among the novel candidate interactors identified in the E2 and TOPBP1 proteomic analysis was nucleolin (NCL), a multifunctional nucleolar protein involved in ribosome biogenesis, chromatin remodeling, DNA repair, replication, and recombination (29, 45–50). To determine whether HPV16 influences NCL RNA expression, we measured NCL mRNA levels in N/Tert-1, N/Tert-1+HPV16, HFK-E6E7, and HFK-HPV16 cells by qRT-PCR (Figure 5A). HPV16-positive cells exhibited significantly higher NCL transcript levels than their donor-matched controls under both monolayer and differentiating conditions (Figure 5A; N/Tert-1+HPV16 vs. N/Tert-1: lanes 2 and 6 vs. lanes 1 and 5; HFK-HPV16 vs. HFK-E6E7: lanes 4 and 8 vs. lanes 3 and 7). As NCL has roles in DNA damage repair; it is critical for DNA double-strand break repair (49, 50), we hypothesize that the upregulation of NCL transcription in HPV16-positive cells is driven by HPV-induced DNA damage during viral genome replication.

**Figure 5:**
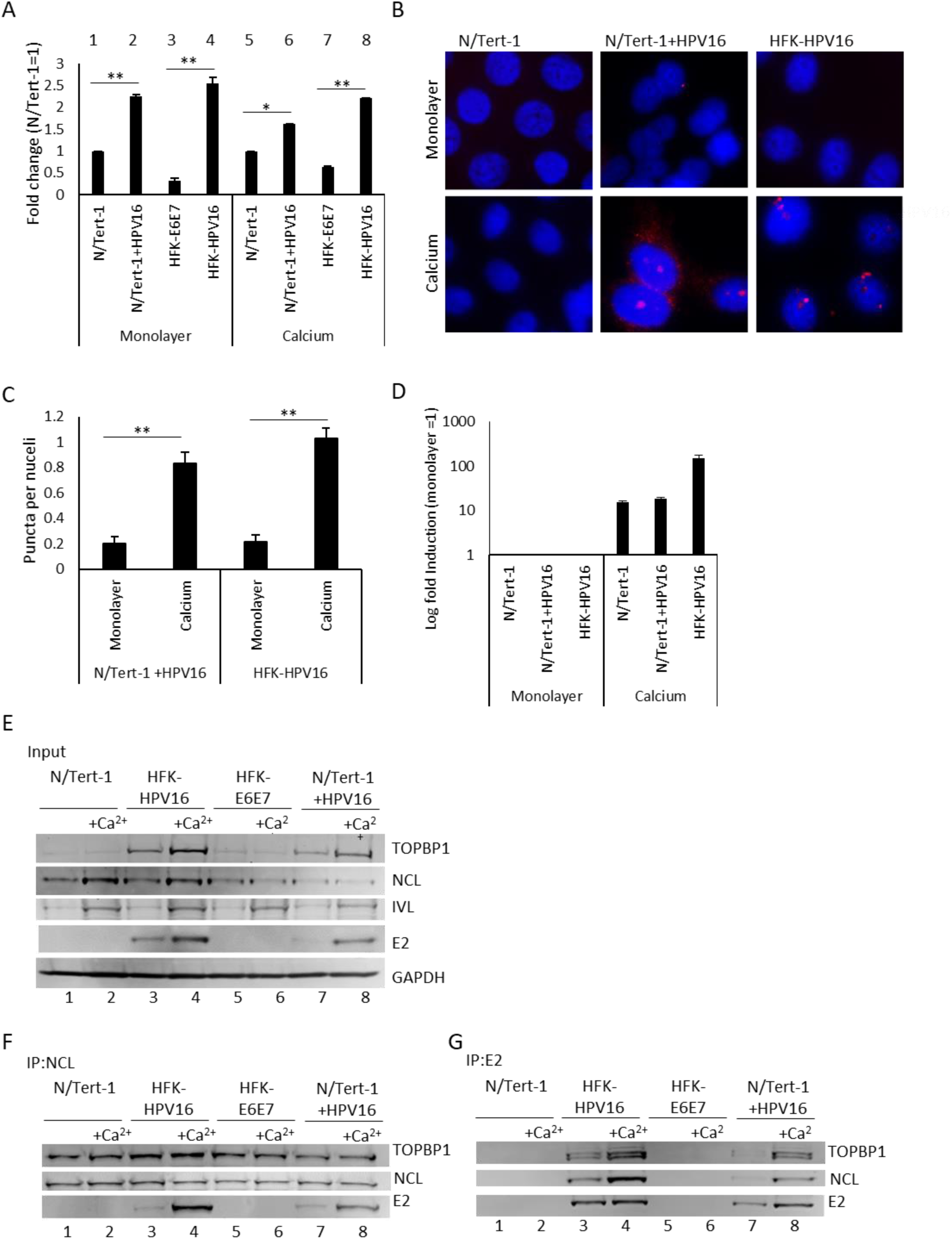
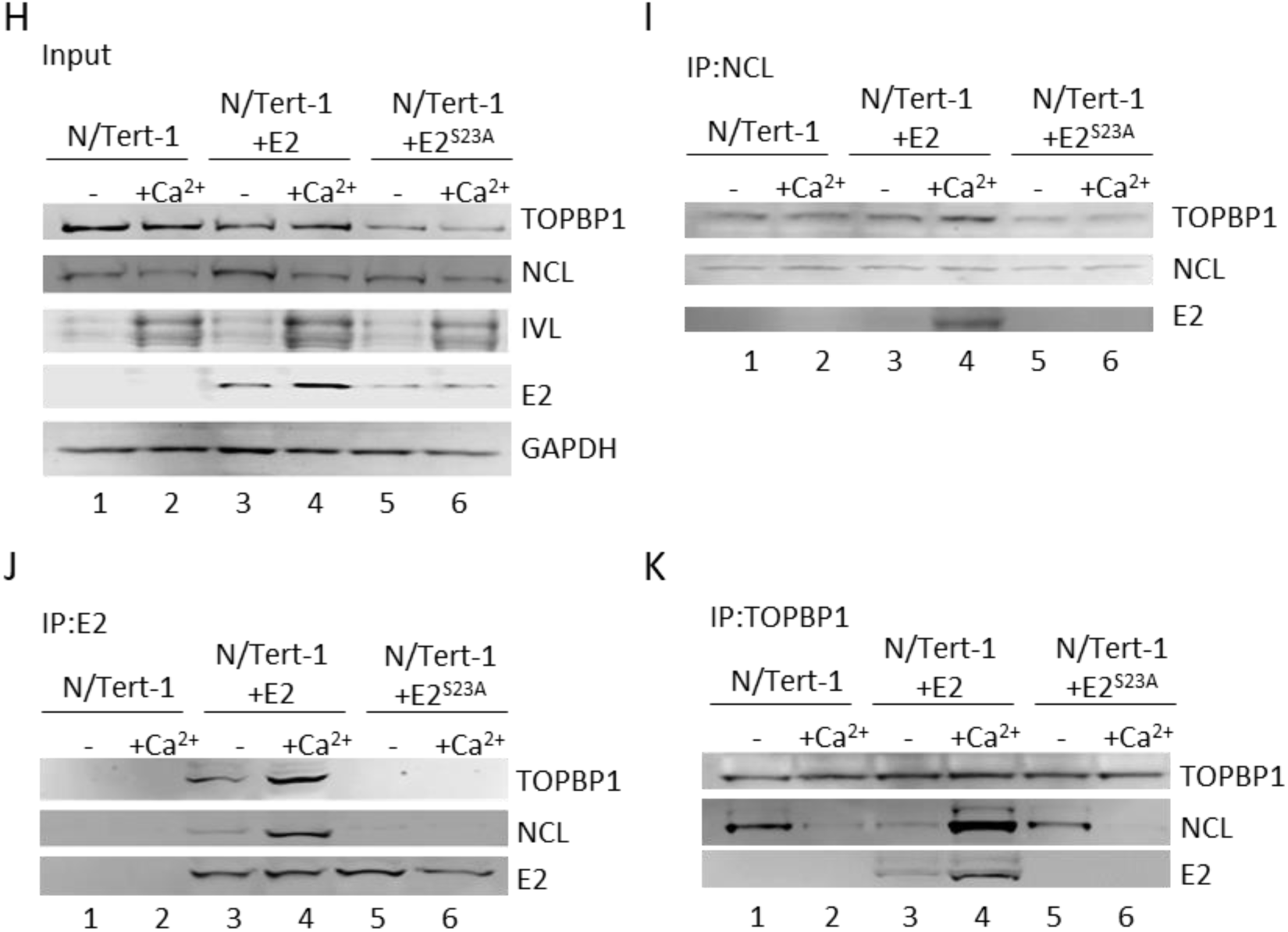
Nucleolin (NCL) interacts with HPV16 E2 in a differentiation- and TOPBP1-dependent manner. A. qRT-PCR analysis of NCL transcript levels in N/Tert-1, N/Tert-1+HPV16, HFK-E6E7 and HFK-HPV16 keratinocytes. Expression is normalized to GAPDH and shown relative to N/Tert-1 controls. Error bars indicate SEM; *P<0.05, **P < 0.01. B. Representative fields of view from proximity ligation assay (PLA) demonstrating *in situ* interaction between E2 and NCL in N/Tert-1+HPV16 and HFK-HPV16 cells under monolayer or differentiating conditions. Each red punctum represents a single protein-protein interaction; nuclei are counterstained with DAPI (blue). N/Tert-1 cells lacking E2 expression serve as a negative control. C. Quantification of PLA foci per nucleus in ≥50 cells per condition. Bars represent mean ± SEM; **P < 0.01. D. qRT-PCR analysis of involucrin (IVL), a marker of keratinocyte differentiation, expression in N/Tert-1, N/Tert-1+HPV16 and HFK-HPV16 cells grown in monolayer or under differentiation conditions. Fold induction is calculated relative to monolayer cultures. E. Input western blot confirming expression of TOPBP1, NCL, IVL and E2 under proliferating and differentiating conditions. GAPDH serves as a loading control. F. Co-immunoprecipitation of NCL and E2. Lysates from growing and differentiating N/Tert-1, N/Tert-1+HPV6, HFK-E6E7 and HFK-HPV16 cells were immunoprecipitated with anti-NCL antibody, followed by western blotting for E2 (B9) and TOPBP1. G. Reciprocal immunoprecipitation of NCL and E2. Lysates from the same cell types and conditions as (F) were immunoprecipitated with anti-E2 antibody, followed by western blotting with NCL and TOPBP1. H. Input western blot confirming expression of TOPBP1, NCL, IVL and E2 in growing and differentiating N/Tert-1, N/Tert-1+E2 and N/Tert-1+E2S23A cells. I. Co-immunoprecipitation of E2 and TOPBP1 with NCL. Lysates from growing and differentiating N/Tert-1, N/Tert-1+E2 and N/Tert-1+E2S23A were immunoprecipitated with anti-NCL antibody, followed by western blotting with E2 and TOPBP1. J. Reciprocal immunoprecipitation of NCL and TOPBP1 with E2. Lysates from the same conditions as (I) were immunoprecipitated with anti-E2 antibody, followed by western blotting with NCL and TOPBP1. K. Reciprocal immunoprecipitation of TOPBP1 and E2 with NCL. Lysates from the same conditions as (I) were immunoprecipitated with anti-TOPBP1 antibody, followed by western blotting with NCL and E2.

To validate the E2-NCL interaction in our model of the HPV16 life cycle, we performed PLA for E2 and NCL in both monolayer and differentiating N/Tert-1+HPV16 and HFK-HPV16, human foreskin keratinocytes immortalized by HPV16 that retain expression of E2 (20, 31, 51). N/Tert-1 cells were utilized as an E2-negative control. While minimal signal was observed in monolayer grown cells, strong nuclear PLA foci were detected in differentiating N/Tert-1+HPV16 and HFK-HPV16 cells (Figure 5B). This signal was quantified computationally, to find that the observed increase in PLA signal in differentiating cells was significant in both N/Tert-1+HPV16 and HFK-HPV16 E2-NCL (Figure 5C). Induction of differentiation was confirmed by qRTPCR for involucrin (IVL); culture in calcium induced expression of the differentiation marker by at least 15-fold IVL (Figure 5D).

To confirm the E2-NCL interaction via an independent approach, reciprocal co-immunoprecipitations (co-IPs) with anti-E2 and anti-NCL antibodies were performed in the same panel of cell lines (N/Tert-1, N/Tert-1+HPV16 and HFK-HPV16), cultured under both monolayer and calcium-induced differentiating conditions (Figure 5E-G). Parental N/Tert-1 and HFKs immortalized by E6 and E7 (HFK-E6E7) served as an E2-negative, donor controls for N/Tert-1+HPV16 and HFK-HPV16 respectively. Input blots (Figure 5E) confirm prior findings that TOPBP1 protein levels are elevated in E2-expressing cells (lanes 3,4,7 and 8) and further increase upon differentiation (lanes 4 and 8)(20, 22). As expected, IVL protein expression increased under differentiating conditions (lanes 2,4,6 and 8), confirming the effectiveness of calcium treatment (52). Reciprocal co-IPs demonstrated that NCL and E2 interact in our model of the HPV16 life cycle (Figure 5F and G). NCL immunoprecipitation pulled down E2 in both HFK-HPV16 and N/Tert-1+HPV16 (Figure 5F, lanes 3,4,7, and 8), and most robustly under differentiating conditions in both cell lines (lanes 4 and 8). E2 IP recovered NCL with similar differentiation dependence (Figure 5G, lanes 4 and 8). This confirms the interaction between E2 and NCL in keratinocytes and indicates that their association is enhanced upon differentiation. Together, the PLA and co-IP results suggests a role for NCL in viral genome amplification, which occurs in mid-layers of epithelia (12).

As NCL was identified as an interactor with both E2 and TOPBP1 in the proteomic screen (Supplementary Table), we next investigated whether the interaction with NCL interacts depends on the presence of both proteins. Co-immunoprecipitation experiments were performed in N/Tert-1 cells stably expressing HPV16 E2 or a TOPBP1-binding deficient mutant (E2^S23A^) (8, 22), under proliferating and calcium-induced differentiating conditions (Figure 5H-K). Parental N/Tert-1 cells served as an E2-negative control. Consistent with previous findings, input blots confirmed that the E2-TOPBP1 interaction stabilizes both proteins (22), as overall E2 and TOPBP1 protein levels were reduced in cells expressing the E2^S23A^ mutant (Figure 5H, lanes 5 and 6). Total NCL expression remain unchanged by either E2 or E2^S23A^ expression, compared to N/Tert-1 or by differentiation (lanes 3-6). Increased IVL expression confirmed calcium-induced differentiation (lanes 2,4, and 6). Immunoprecipitation with anti-NCL antibody (Figure 5I) confirmed E2 pulldown in calcium treated cells (Figure 5I, lane 4), whereas no E2 was pulled down in E2^S23A^ -expressing cells (lanes 5 and 6). Together, these results demonstrate that NCL associates with E2 in a differentiation- and TOPBP1-dependent manner. Consistent with these findings, E2 immunoprecipitations showed that wild-type E2 efficiently co-immunoprecipitated both TOPBP1 and NCL, whereas E2^S23A^ failed to do so, indicating that NCL association with E2 requires TOPBP1 binding (Figure 5J, lanes 3 and 4 compared to 5 and 6). Using the E2 antibody, the E2:NCL interaction was seen in monolayer and was further enhanced under differentiating conditions (lanes 3 and 4). Reciprocal immunoprecipitation with anti-TOPBP1 antibody showed that TOPBP1 and NCL interact in the absence of E2 (Figure 5K), with a stronger association observed in monolayer cultures (Figure 5K, lanes 1 and 2). However, in the presence of E2, differentiation strengthened the TOPBP1-NCL association (lanes 3 and 4) whereas E2^S23A^ expression abrogated this effect (lanes 5 and 6).

Taken together these findings indicate that E2 promotes or stabilizes a differentiation-dependent complex between TOPBP1 and NCL, and that the E2-NCL interaction requires TOPBP1 binding. NCL may function as part of an E2–TOPBP1–NCL complex involved in amplification stages of the HPV life cycle.

#### NCL supports viral genome maintenance

Having established that NCL interacts with E2 in a differentiation- and TOPBP1-dependent manner, we next sought to determine the functional significance of this interaction in the HPV16 life cycle. Given NCL’s roles in chromatin organization, replication, and DNA damage responses, and its differentiation-dependent association with E2 and TOPBP1, we hypothesized that NCL contributes to viral genome maintenance or amplification. To test this, we examined the effects of NCL depletion on viral genome replication in HPV16-positive keratinocytes (Figure 6). We performed siRNA-mediated knockdown of NCL in HPV16-episome containing HFKs cultured under proliferating (monolayer) and differentiating (high-calcium) conditions. Knockdown efficiency was confirmed by qRT-PCR, which showed >60% reduction in NCL transcript levels relative to non-targeting controls in both N/Tert-1+HPV16 and HFK-HPV16 cells (Figure 6A), demonstrating reduced NCL protein abundance. Differentiation was verified by increased IVL expression quantified by qRT-PCR (Figure 6B). To assess the impact of NCL depletion on viral genome status, we employed an exonuclease V (ExoV) assay, which selectively digests linear DNA but spares circular episomes; a robust alternative method to Southern blotting that enables sensitive detection with lower input requirements and greater quantitative power (5, 53). In differentiating HFK-HPV16 cells, depletion of NCL led to a reduction in ExoV-resistant HPV16 DNA, indicating a loss of episomal genomes during the amplification phase of the viral life cycle (Figure 6C). Integration, measured by degradation of linear E2 and E6 DNA following ExoV treatment, was significantly increased upon NCL knockdown with two independent siRNAs compared with control siRNA (*p*<0.01 for both E2 and E6 siRNA1 and E2 siRNA2, *p*<0.05 E6 siRNA2). By contrast, equivalent NCL knockdown in monolayer HFK-HPV16 cells did not significantly alter episomal or integrated viral DNA levels, demonstrating that NCL is specifically required for episome maintenance in differentiating keratinocytes. Calcium-induced differentiation alone modestly increased viral integration; however, this effect reached statistical significance only when measured with E6 primers (*p*<0.05), not with E2. Given the role of NCL in chromatin remodeling and nucleolar organization, these results suggest that the E2-NCL association is important for stabilizing viral episomes during genome amplification and/or for protecting viral DNA from nucleolytic degradation (29, 48).

**Figure 6:**
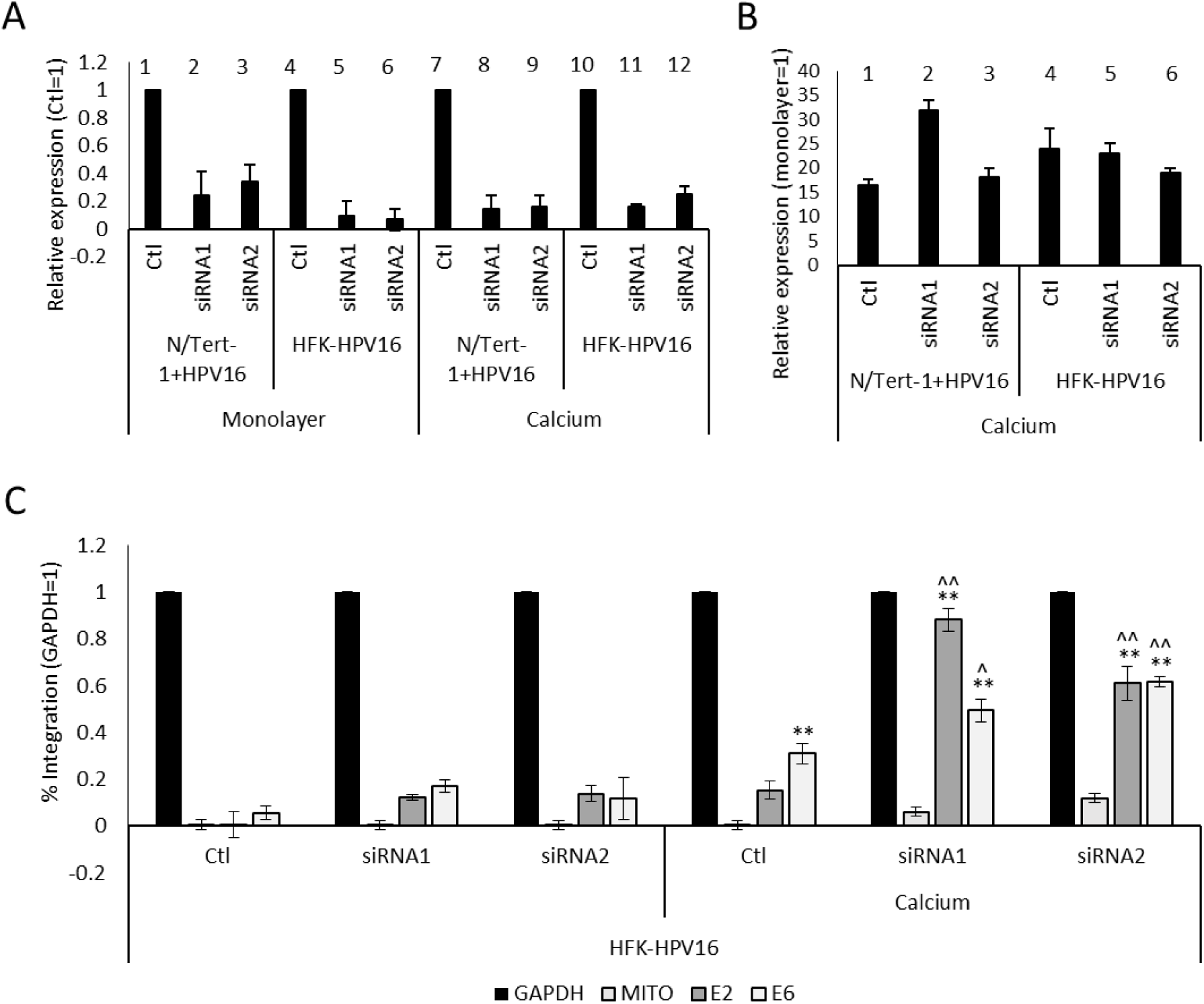
NCL supports viral genome maintenance. A. Confirmation of siRNA-mediated knockdown of NCL in N/Tert-1, N/Tert-1+HPV16 and HFK-HPV16 cells by qRT-PCR, showing efficient depletion of NCL transcripts in both monolayer and differentiating conditions, by two separate siRNAs, compared to non-targeting control (Ctl). B. Quantification of IVL, a marker of keratinocyte differentiation, expression in NCL siRNA-treated N/Tert-1, N/Tert-1+HPV16 and HFK-HPV16 in siRNA treated cells. C. Exonuclease V (ExoV) digestion assay HFK-HPV16 cells. ExoV selectively degrades linear DNA but not circular episomal DNA. Knockdown of NCL reduced the proportion of ExoV-resistant HPV16 genomes in calcium treated cells, indicating loss of episomal viral DNA during amplification of the viral genome. Bars represent the mean of three biological replicates; error bars indicate standard error of the mean (SEM). Statistical significance was assessed using unpaired t-tests; **p < 0.01 compared to monolayer equivalent, ^p<0.05, ^^p<0.01 compared to control siRNA.

## Discussion

In this study, we employed TurboID proximity proteomics to map the HPV16 E2 interactome in human keratinocytes under both basal and differentiating conditions. By using an inducible expression system in a physiologically relevant host cell, our approach provides a proteomic perspective that both confirms and expands on prior studies (17, 30). Previous work has demonstrated that HPV16 E2 recruits host repair proteins such as BRD4 and TOPBP1 to viral replication centers, facilitating viral genome maintenance (20–22, 24, 40). Our analysis reinforces this model while also revealing an enrichment of pathways linked to chromatin organization and RNA metabolism. Notably, the identification of spliceosome- and translation-related factors suggests that E2 may also influence post-transcriptional regulation of viral and host gene expression, consistent with earlier reports linking E2 to transcriptional control (8, 54). Altogether, this analysis reveals that HPV16 E2 engages in a broad network of host factors spanning chromatin organization, transcriptional regulation, RNA splicing, and translation.

Direct comparison of the E2 and TOPBP1 interactomes revealed significant overlap, underscoring the functional partnership of these two proteins. In basal cells, the E2-TOPBP1 interaction likely recruits host DNA replication and repair complexes, consistent with the established role of the E2-TOPBP1 complex in HPV replication (20, 21). In differentiating cells, TOPBP1-associated networks included translation- and RNA processing–related factors, suggesting that HPV16 modifies TOPBP1 complexes to redirect host machinery toward productive replication. We propose that E2 hijacks this versatility to manipulateTOPBP1 complexes in a differentiation-dependent manner, balancing genome maintenance in basal layers with productive amplification in suprabasal cells.

The HPV life cycle is intimately linked to epithelial differentiation, with genome maintenance in basal cells and genome amplification occurring in suprabasal layers (12). Nucleolin (NCL) emerged from this study as a differentiation- and TOPBP1-dependent partner of E2. Notably, TOPBP1 binding is crucial for the E2–NCL interaction, and E2 expression enhances the TOPBP1–NCL association, suggesting a cooperative mechanism. Beyond its role in nucleolar organization and chromatin remodeling (44,45,47), NCL participates in the DNA damage response by facilitating repair factor recruitment and promoting genome stability (46, 48–50). Depletion of NCL compromised the stability of episomal HPV16 genomes during differentiation, leading to a significant reduction in ExoV-resistant viral DNA and increased degradation of linear E2 and E6 fragments, consistent with enhanced viral integration. This effect was specific to differentiating cells and not observed in monolayer cultures, highlighting NCL’s role in maintaining episomal genome stability during the amplification phase of the viral life cycle. These observations align with accumulating evidence that E2 coordinates with host DNA damage response (DDR) factors to preserve viral episomes and regulate replication (11, 20, 21, 55, 56). Disruption of these interactions, such as by knockdown of SAMHD1 or SMARCAL1, similarly promotes viral genome integration, underscoring the importance of E2-DDR factor recruitment in episomal stability (55, 56). Together, these findings suggest that E2 exploits NCL, in concert with TOPBP1, to coordinate DNA damage signaling and chromatin reorganization at viral replication centers, particularly during differentiation-dependent genome amplification. Furthermore, E2 and TOPBP1 have recently been shown to localize to nuclear condensates (17), indicating that NCL may contribute to the recruitment or stabilization of E2 within these specialized subnuclear domains. Future studies will investigate the mechanistic basis of the E2– TOPBP1–NCL complex in nucleolar function and viral genome amplification, as well as its potential role in nuclear condensate dynamics.

Importantly, NCL represents a potential therapeutic target. Overexpression of NCL correlates with poor prognosis in multiple cancers, including NSCLC and AML, and NCL-targeting strategies are under active investigation (48–52). Given that E2 expression persists in a subset of HPV-positive head and neck cancers (4, 5), disruption of E2–TOPBP1-NCL interactions could impair both episome stability and tumor maintenance. Targeting this complex may therefore provide dual benefits: interfering with viral genome maintenance and potentially disrupting oncogenic processes that rely on persistent E2 expression. These findings highlight the E2– TOPBP1–NCL interaction as a promising candidate for therapeutic intervention in HPV-driven malignancies.

In summary, our work demonstrates that HPV16 E2 engages dynamic, differentiation-dependent host interaction networks in keratinocytes. By comparing the interactomes of E2 and TOPBP1, we reveal both shared and unique host partners that likely contribute to viral genome maintenance, productive replication, and persistence. We propose a model in which E2 coordinates distinct sets of host interactions across the epithelial life cycle, balancing genome maintenance in basal cells with amplification in suprabasal cells. The identification of NCL as a differentiation-dependent partner, whose interaction with E2 is facilitated by TOPBP1, highlights a novel complex important for episomal genome stability. These findings provide new insight into HPV biology and underscore host pathways that may be exploited to interfere with the viral life cycle.

## Materials and Methods

### Cell culture and differentiation

Human telomerase-immortalized keratinocytes (N/Tert-1) and N/Tert-1 cells stably harboring HPV16 episomes (N/Tert-1+HPV16) were cultured in keratinocyte serum-free medium (kSFM, Gibco) supplemented with 0.2 ng/ml EGF, 30 μg/ml bovine pituitary extract, 0.3 mM calcium chloride and 7.5 µM hygromycin. N/Tert-1 cells stably expressing wild-type E2 and E2^S23A^ were generated and cultured as previously described (8, 22). C33A cervical carcinoma cells were maintained in DMEM (Gibco) supplemented with 10% fetal bovine serum (FBS). HPV16- and E6E7-immortalized human foreskin keratinocytes (HFKs), previously described (51), were cultured in DermaLife-K Complete Medium (Lifeline Cell Technology). All HPV16-positive cells were co-cultured with mitomycin C–inactivated J2 fibroblasts. All cell lines were routinely tested for mycoplasma. To induce differentiation, cells were cultured to 70% confluency, incubated overnight in low-calcium media (M154 CF; Invitrogen), and then switched to 1.5 mM CaCl_2_-containing medium for 72 h.

### Plasmids, lentiviral transduction and siRNA

The HPV16 E2 and TOPBP1 open reading frames were cloned separately into the pCW57.1_V5_TurboID_FLAG_Nt vector (Addgene #118055; a gift from the RESOLUTE Consortium and Giulio Superti-Furga) by Genscript, generating doxycycline-inducible constructs expressing N-terminal TurboID-3×FLAG–tagged proteins with a C-terminal V5 epitope. Lentiviral particles were produced in HEK293T cells using standard third-generation packaging plasmids. Target cells were transduced in the presence of 8 μg/ml polybrene and selected with 1 μg/ml (N/Tert-1) or 10 μg/ml (C33a) blasticidin. MISSION® esiRNA targeting NCL and control siRNA were purchased from SigmaAldrich. siRNA targeting the TOPBP1 3’UTR was purchased from Thermo Fisher. Cells were transfected with 10 uM siRNA and harvested 48 h later for DNA, RNA, and protein analysis.

### Induction and biotin labelling

TurboID-tagged E2 or TOPBP1 was induced with 5 μM doxycycline for 48 h. For biotin labeling, cells were supplemented with 50 μM biotin (Sigma) for the final 6 h of induction. Vehicle-treated cells (ethanol or PBS) served as controls.

### Transient DNA Replication Assay

Replication assays were performed in C33A cells co-transfected with plasmids encoding the HPV16 origin (pOri), E1, and wild-type HPV16 E2. TurboID expression was induced with 5 μM doxycycline or vehicle 24 h post-transfection. Low-molecular-weight DNA was harvested 72 h post-transfection using the Hirt method. DNA was digested with DpnI to remove input plasmid and quantified by qPCR using primers specific to the HPV16 origin as described previously(28). pOri primers: Fwd 5′-ATCGGTTGAACCGAAACCG-3′; Rev 5′-TAACTTCTGGGTCGCTCCTG-3′.

### Luciferase Transcription Assay

N/Tert-1 cells stably expressing TurboID constructs were pretreated with 5 μM doxycycline or vehicle for 24 h, followed by transfection with either pTKE2 or LCR luciferase reporter plasmids using Lipofectamine 3000. Luciferase activity was measured 48 h later using the Dual-Luciferase Reporter Assay System (Promega) and normalized to total protein concentration.

### Western Blotting

Cells were lysed in T-PER buffer (Thermo Scientific) supplemented with protease and phosphatase inhibitors (Roche, MilliporeSigma). Lysates were clarified by centrifugation at 18,400 × g for 20 min at 4°C, and protein concentration was determined using the Bio-Rad Protein Assay. Equal amounts (50 μg) of protein were resolved by SDS–PAGE (Novex 4–12% Tris-glycine gels; Invitrogen) and transferred to nitrocellulose membranes (Bio-Rad) overnight at 30 V. Membranes were blocked in Odyssey PBS blocking buffer (LI-COR, diluted 1:1 in PBS) for 1 h at room temperature and incubated overnight at 4°C with primary antibodies. After washing in PBS–0.1% Tween-20, membranes were probed with IRDye-conjugated secondary antibodies (LI-COR) at 1:10,000. Signal was visualized on an Odyssey CLx Imaging System and quantified in ImageJ. GAPDH (Santa Cruz 47724) was used as a loading control.

Primary antibodies: TOPBP1 (ThermoFisher A300-111A), BirA (ThermoFisher PA5-80250), Nucleolin (ThermoFisher PA3-16875), ATR (Cell Signaling 2790), ATM (Cell Signaling 2873), phospho-ATR (Cell Signaling 2853), phospho-ATM (ThermoFisher MA1-46069), CHK1 (Cell Signaling 2360), CHK2 (Cell Signaling 2622), phospho-CHK1 (Cell Signaling 2197), phospho-CHK2 (Cell Signaling 2197), HPV16 E2 (clone B9)(57) and γH2AX (phospho-S139; Cell Signaling 9718), NCL (life tech; PA3-16875), IVL (SCBT; sc-21748).

### Immunoprecipitation

Cell lysates were prepared as above. 250 μg protein was incubated with pre-washed streptavidin beads (Pierce, Thermo Scientific) overnight at 4°C with rotation. Beads were washed 3× in lysis buffer, resuspended in Laemmli sample buffer, denatured at 95°C, and analyzed by SDS–PAGE and immunoblotting. For proteomics, beads were washed once in nuclease-free water before processing.

### Proteomics and bioinformatics

Biotinylated proteins enriched on streptavidin beads were processed for LC–MS/MS using the PreOmics iST kit (PreOmics GmbH) following manufacturer instructions. Peptides were analyzed by LC–MS/MS and searched against the UniProt human proteome supplemented with HPV16 sequences using Sequest HT in Proteome Discoverer v3.0 (Thermo). Protein identifications were filtered at a false discovery rate (FDR) of <0.01, and quantification was based on peptide intensities normalized to total abundance. Venn diagrams were generated using InteractiVenn(58). Gene ontology analysis was performed using ShinyGO v0.82(59) with a human background, a minimum pathway size of 2, and FDR <0.05.

### qRTPCR

RNA was isolated with the SV Total RNA Isolation kit (Promega) and reverse-transcribed with the High-Capacity cDNA Kit (Invitrogen). qPCR was performed with PowerUp SYBR Green Master Mix (Applied Biosystems) on a 7500 Fast system. Expression was normalized to GAPDH using the 2^-ΔΔCT method. Primers: GAPDH (Fwd 5′-GGAGCGAGATCCCTCCAAAAT-3′; Rev 5′-GGCTGTTGTCATACTTCTCATGG-3′), IVL (Fwd 5′-TCCTCCAGTCAATACCCATCAG-3′; Rev 5′-CAGCAGTCATGTGCTTTTCCT-3′), BirA (Fwd 5′-TCCTGGCTAATGGCGAGTTC-3′; Rev 5′-TAGGCTCGGGCAGAGAGTAG-3′), NCL (Fwd 5′-GGTGGTCGTTTCCCCAACAAA-3′; Rev 5′-GGTGGTCGTTTCCCCAACAAA-3′).

### Immunofluorescence (IF)

Cells grown on acid-etched glass coverslips were fixed in ice-cold methanol for 10 min, permeabilized with 0.2% Triton X-100/PBS for 15 min, blocked in 5% BSA, and incubated with primary antibodies (E2 TVG261, TOPBP1, NCL, BirA). Alexa Fluor 488– or 594–conjugated secondary antibodies (Molecular Probes) were used for detection. Nuclei were counterstained with DAPI and mounted in Vectashield (ThermoFisher). Images were acquired on a Keyence fluorescence microscope.

### Proximity Ligation Assay (PLA)

Protein–protein interactions were visualized using the Duolink® In Situ Red Kit (MilliporeSigma) according to the manufacturer’s protocol. Cells were incubated with primary antibodies from different species, followed by PLUS and MINUS probes, ligation, and rolling circle amplification. Nuclei were counterstained with DAPI. Negative controls omitting one or both primary antibodies were included.

### Exonuclease V Assay

Viral genome status was determined using Exonuclease V digestion followed by qPCR, as described(53). Genomic DNA (20 ng) was treated with Exonuclease V (NEB) for 1 h at 37°C and heat-inactivated. Treated and untreated DNA was analyzed by qPCR using primers specific for HPV16 E6, E2, human mitochondrial DNA (MITO), and GAPDH. Primers: HPV16 E6 Fwd: 5’-TTGCTTTTCGGGATTTATGC-3’ Rev: 5’-CAGGACACAGTGGCTTTTGA-3’, HPV16 E2 Fwd: 5’-TGGAAGTGCAGTTTGATGGA-3’ Rev: 5’-CCGCATGAACTTCCCATACT-3’, MITO Fwd: 5’-CAGGAGTAGGAGAGAGGGAGGTAAG-3’ Rev: 5’-TACCCATCATAATCGGAGGCTTTGG -3’, GAPDH Fwd: 5’-GGAGCGAGATCCCTCCAAAAT-3’ Rev: 5’-GGCTGTTGTCATACTTCTCATGG-3’

### Statistical analysis

All experiments were performed in ≥3 biological replicates. Data are presented as mean ± standard error of the mean (SEM). Comparisons between two groups were assessed by unpaired Student’s t test. p < 0.05 was considered statistically significant. Proteomic significance was determined using a 1% FDR and SAINT scoring where applicable.

## Acknowledgements

Services and products in support of the research project were generated by the VCU Massey Comprehensive Cancer Center Proteomics Shared Resource, supported, in part, with funding from NIH-NCI Cancer Center Support Grant P30 CA016059. This work was supported by R01DE029471 (IMM) and R21AI178143 (IMM). Figure 1A and B were created with Biorender.

## References

1. Gribb JP, Wheelock JH, Park ES. 2023. Human Papilloma Virus (HPV) and the Current State of Oropharyngeal Cancer Prevention and Treatment. Del J Public Health 9:26–28.

2. Rettig EM, Sethi RKV. 2021. Cancer of the Oropharynx and the Association with Human Papillomavirus. Hematol Oncol Clin North Am 35:913–931.

3. Malik S, Sah R, Muhammad K, Waheed Y. 2023. Tracking HPV Infection, Associated Cancer Development, and Recent Treatment Efforts—A Comprehensive Review. Vaccines 11:102.

4. Koneva LA, Zhang Y, Virani S, Hall PB, McHugh JB, Chepeha DB, Wolf GT, Carey TE, Rozek LS, Sartor MA. 2018. HPV Integration in HNSCC Correlates with Survival Outcomes, Immune Response Signatures, and Candidate Drivers. Mol Cancer Res 16:90– 102.

5. James CD, Otoa RO, Youssef AH, Fontan CT, Sannigrahi MK, Windle B, Basu D, Morgan IM. 2024. HPV16 genome structure analysis in oropharyngeal cancer PDXs identifies tumors with integrated and episomal genomes. Tumour Virus Res 18:200285.

6. Nulton TJ, Olex AL, Dozmorov M, Morgan IM, Windle B. 2017. Analysis of The Cancer Genome Atlas sequencing data reveals novel properties of the human papillomavirus 16 genome in head and neck squamous cell carcinoma. Oncotarget 8:17684–17699.

7. Anayannis NV, Schlecht NF, Ben-Dayan M, Smith RV, Belbin TJ, Ow TJ, Blakaj DM, Burk RD, Leonard SM, Woodman CB, Parish JL, Prystowsky MB. 2018. Association of an intact E2 gene with higher HPV viral load, higher viral oncogene expression, and improved clinical outcome in HPV16 positive head and neck squamous cell carcinoma. PLOS ONE 13:e0191581.

8. Evans MR, James CD, Bristol ML, Nulton TJ, Wang X, Kaur N, White EA, Windle B, Morgan IM. 2019. Human Papillomavirus 16 E2 Regulates Keratinocyte Gene Expression Relevant to Cancer and the Viral Life Cycle. J Virol 93:e01941–18.

9. Morgan IM. 2025. The functions of papillomavirus E2 proteins. Virology 603:110387.

10. McBride AA. 2013. The Papillomavirus E2 proteins. Virology 445:57–79.

11. Prabhakar AT, James CD, Nguyen AQ, Bridy PV, Wang X, Zoldork RJ, Roe JD, Witt A, Panidepu SS, Bristol ML, Dalton K, Wieland A, Diab A, Faber AC, Androphy EJ, Basu D, Lambert P, Spurgeon ME, Morgan IM. 2025. E2 displacement of CIP2A from TOPBP1 activates the DNA damage response during papillomavirus life cycles. BioRxiv 10.1101/2025.10.15.682653.

12. Doorbar J, Quint W, Banks L, Bravo IG, Stoler M, Broker TR, Stanley MA. 2012. The Biology and Life-Cycle of Human Papillomaviruses. Vaccine 30:F55–F70.

13. Baxter MK, McPhillips MG, Ozato K, McBride AA. 2005. The Mitotic Chromosome Binding Activity of the Papillomavirus E2 Protein Correlates with Interaction with the Cellular Chromosomal Protein, Brd4. J Virol 79:4806–4818.

14. DeSmet M, Jose L, Isaq N, Androphy EJ. 2019. Phosphorylation of a Conserved Tyrosine in the Papillomavirus E2 Protein Regulates Brd4 Binding and Viral Replication. J Virol 93:e01801–18.

15. Abbate EA, Voitenleitner C, Botchan MR. 2006. Structure of the Papillomavirus DNA-Tethering Complex E2:Brd4 and a Peptide that Ablates HPV Chromosomal Association. Mol Cell 24:877–889.

16. Krüppel U, Müller-Schiffmann A, Baldus SE, Smola-Hess S, Steger G. 2008. E2 and the co-activator p300 can cooperate in activation of the human papillomavirus type 16 early promoter. Virology 377:151–159.

17. Peng Y-C, Breiding DE, Sverdrup F, Richard J, Androphy EJ. 2000. AMF-1/Gps2 Binds p300 and Enhances Its Interaction with Papillomavirus E2 Proteins. J Virol 74:5872–5879.

18. Harris L, McFarlane-Majeed L, Campos-León K, Roberts S, Parish JL. 2017. The Cellular DNA Helicase ChlR1 Regulates Chromatin and Nuclear Matrix Attachment of the Human Papillomavirus 16 E2 Protein and High-Copy-Number Viral Genome Establishment. J Virol 91:e01853–16.

19. Boner W, Morgan IM. 2002. Novel cellular interacting partners of the human papillomavirus 16 transcription/replication factor E2. Virus Res 90:113–118.

20. Prabhakar AT, James CD, Fontan CT, Otoa R, Wang X, Bristol ML, Hill RD, Dubey A, Morgan IM. 2023. Human Papillomavirus 16 E2 Interaction with TopBP1 Is Required for E2 and Viral Genome Stability during the Viral Life Cycle. J Virol 97:e00063–23.

21. Prabhakar AT, James CD, Youssef AH, Hossain RA, Hill RD, Bristol ML, Wang X, Dubey A, Karimi E, Morgan IM. 2024. A human papillomavirus 16 E2-TopBP1 dependent SIRT1-p300 acetylation switch regulates mitotic viral and human protein levels and activates the DNA damage response. mBio 15:e0067624.

22. Prabhakar AT, James CD, Das D, Otoa R, Day M, Burgner J, Fontan CT, Wang X, Glass SH, Wieland A, Donaldson MM, Bristol ML, Li R, Oliver AW, Pearl LH, Smith BO, Morgan IM. 2021. CK2 Phosphorylation of Human Papillomavirus 16 E2 on Serine 23 Promotes Interaction with TopBP1 and Is Critical for E2 Interaction with Mitotic Chromatin and the Viral Life Cycle. mBio 12:e0116321.

23. Cho KF, Branon TC, Udeshi ND, Myers SA, Carr SA, Ting AY. 2020. Proximity labeling in mammalian cells with TurboID and split-TurboID. Nat Protoc 15:3971–3999.

24. Frattini C, Promonet A, Alghoul E, Vidal-Eychenie S, Lamarque M, Blanchard M-P, Urbach S, Basbous J, Constantinou A. 2021. TopBP1 assembles nuclear condensates to switch on ATR signaling. Mol Cell 81:1231–1245.e8.

25. Bagge J, Oestergaard VH, Lisby M. 2021. Functions of TopBP1 in preserving genome integrity during mitosis. Semin Cell Dev Biol 113:57–64.

26. Bang SW, Ko MJ, Kang S, Kim GS, Kang D, Lee J, Hwang DS. 2011. Human TopBP1 localization to the mitotic centrosome mediates mitotic progression. Exp Cell Res 317:994– 1004.

27. Donaldson MM, Mackintosh LJ, Bodily JM, Dornan ES, Laimins LA, Morgan IM. 2012. An Interaction between Human Papillomavirus 16 E2 and TopBP1 Is Required for Optimum Viral DNA Replication and Episomal Genome Establishment. J Virol 86:12806–12815.

28. Boner W, Taylor ER, Tsirimonaki E, Yamane K, Campo MS, Morgan IM. 2002. A Functional Interaction between the Human Papillomavirus 16 Transcription/Replication Factor E2 and the DNA Damage Response Protein TopBP1. J Biol Chem 277:22297– 22303.

29. Mongelard F, Bouvet P. 2007. Nucleolin: a multiFACeTed protein. Trends Cell Biol 17:80– 86.

30. Branon TC, Bosch JA, Sanchez AD, Udeshi ND, Svinkina T, Carr SA, Feldman JL, Perrimon N, Ting AY. 2018. Efficient proximity labeling in living cells and organisms with TurboID. Nat Biotechnol 36:880–887.

31. Evans MR, James CD, Loughran O, Nulton TJ, Wang X, Bristol ML, Windle B, Morgan IM. 2017. A keratinocyte life cycle model identifies novel host genome regulation by human papillomavirus 16 relevant to HPV positive head and neck cancer. Oncotarget 8:81892– 81909.

32. Guo J, Guo S, Lu S, Gong J, Wang L, Ding L, Chen Q, Liu W. 2023. The development of proximity labeling technology and its applications in mammals, plants, and microorganisms. Cell Commun Signal 21:269.

33. Vance KW, Campo MS, Morgan IM. 1999. An Enhanced Epithelial Response of a Papillomavirus Promoter to Transcriptional Activators. J Biol Chem 274:27839–27844.

34. Vance KW, Campo MS, Morgan IM. 2001. A Novel Silencer Element in the Bovine Papillomavirus Type 4 Promoter Represses the Transcriptional Response to Papillomavirus E2 Protein. J Virol 75:2829–2838.

35. Bouvard V, Storey A, Pim D, Banks L. 1994. Characterization of the human papillomavirus E2 protein: evidence of trans-activation and trans-repression in cervical keratinocytes. EMBO J 13:5451–5459.

36. Demeret C, Desaintes C, Yaniv M, Thierry F. 1997. Different mechanisms contribute to the E2-mediated transcriptional repression of human papillomavirus type 18 viral oncogenes. J Virol 71:9343–9349.

37. Romanczuk H, Thierry F, Howley PM. 1990. Mutational analysis of cis elements involved in E2 modulation of human papillomavirus type 16 P97 and type 18 P105 promoters. J Virol 64:2849–2859.

38. Moody CA, Laimins LA. 2009. Human Papillomaviruses Activate the ATM DNA Damage Pathway for Viral Genome Amplification upon Differentiation. PLoS Pathog 5:e1000605.

39. Muller M. 2012. The HPV E2-Host Protein-Protein Interactions: A Complex Hijacking of the Cellular Network. Open Virol J 6:173–189.

40. Yigitliler A, Renner J, Simon C, Schneider M, Stubenrauch F, Iftner T. 2021. BRD4S Interacts with Viral E2 Protein To Limit Human Papillomavirus Late Transcription. J Virol 95:e02032–20.

41. Jang MK, Anderson DE, Van Doorslaer K, McBride AA. 2015. A proteomic approach to discover and compare interacting partners of papillomavirus E2 proteins from diverse phylogenetic groups. PROTEOMICS 15:2038–2050.

42. Cescutti R, Negrini S, Kohzaki M, Halazonetis TD. 2010. TopBP1 functions with 53BP1 in the G1 DNA damage checkpoint. EMBO J 29:3723–3732.

43. Leimbacher P-A, Jones SE, Shorrocks A-MK, De Marco Zompit M, Day M, Blaauwendraad J, Bundschuh D, Bonham S, Fischer R, Fink D, Kessler BM, Oliver AW, Pearl LH, Blackford AN, Stucki M. 2019. MDC1 Interacts with TOPBP1 to Maintain Chromosomal Stability during Mitosis. Mol Cell 74:571–583.e8.

44. Maser RS, Monsen KJ, Nelms BE, Petrini JHJ. 1997. hMre11 and hRad50 Nuclear Foci Are Induced During the Normal Cellular Response to DNA Double-Strand Breaks †. Mol Cell Biol 17:6087–6096.

45. Zhang X, Xiao S, Rameau RD, Devany E, Nadeem Z, Caglar E, Ng K, Kleiman FE, Saxena A. 2018. Nucleolin phosphorylation regulates PARN deadenylase activity during cellular stress response. RNA Biol 15:251–260.

46. Prakash K, Satishkartik S, Ramalingam S, Gangadaran P, Gnanavel S, Aruljothi KN. 2025. Investigating the multifaceted role of nucleolin in cellular function and Cancer: Structure, Regulation, and therapeutic implications. Gene 957:149479.

47. Cong R, Das S, Ugrinova I, Kumar S, Mongelard F, Wong J, Bouvet P. 2012. Interaction of nucleolin with ribosomal RNA genes and its role in RNA polymerase I transcription. Nucleic Acids Res 40:9441–9454.

48. Tonello F, Massimino ML, Peggion C. 2022. Nucleolin: a cell portal for viruses, bacteria, and toxins. Cell Mol Life Sci 79:271.

49. Scott DD, Oeffinger M. 2016. Nucleolin and nucleophosmin: nucleolar proteins with multiple functions in DNA repair. Biochem Cell Biol 94:419–432.

50. Goldstein M, Derheimer FA, Tait-Mulder J, Kastan MB. 2013. Nucleolin mediates nucleosome disruption critical for DNA double-strand break repair. Proc Natl Acad Sci 110:16874–16879.

51. Fontan CT, James CD, Prabhakar AT, Bristol ML, Otoa R, Wang X, Karimi E, Rajagopalan P, Basu D, Morgan IM. 2022. A Critical Role for p53 during the HPV16 Life Cycle. Microbiol Spectr 10:e00681–22.

52. Watt FM. 1983. Involucrin and Other Markers of Keratinocyte Terminal Differentiation. J Invest Dermatol 81:S100–S103.

53. Myers JE, Guidry JT, Scott ML, Zwolinska K, Raikhy G, Prasai K, Bienkowska-Haba M, Bodily JM, Sapp MJ, Scott RS. 2019. Detecting episomal or integrated human papillomavirus 16 DNA using an exonuclease V-qPCR-based assay. Virology 537:149– 156.

54. Gauson EJ, Windle B, Donaldson MM, Caffarel MM, Dornan ES, Coleman N, Herzyk P, Henderson SC, Wang X, Morgan IM. 2014. Regulation of human genome expression and RNA splicing by human papillomavirus 16 E2 protein. Virology 468–470:10–18.

55. James CD, Youssef A, Prabhakar AT, Otoa R, Roe JD, Witt A, Lewis RL, Bristol ML, Wang X, Zhang K, Li R, Morgan IM. 2024. Human papillomavirus 16 replication converts SAMHD1 into a homologous recombination factor and promotes its recruitment to replicating viral DNA. J Virol 98:e0082624.

56. James CD, Youssef AH, Roe JD, Cappiello F, Aiello FA, Perdichizzi B, Lewis RL, Witt A, Prabhakar AT, Wang X, Bristol ML, Pichierri P, Morgan IM. 2025. HPV16 recruitment of SMARCAL1 to viral and host replication forks is required for the viral life cycle. Microbiology 10.1101/2025.07.22.666081.

57. Wieland A, Patel MR, Cardenas MA, Eberhardt CS, Hudson WH, Obeng RC, Griffith CC, Wang X, Chen ZG, Kissick HT, Saba NF, Ahmed R. 2021. Defining HPV-specific B cell responses in patients with head and neck cancer. Nature 597:274–278.

58. Heberle H, Meirelles GV, Da Silva FR, Telles GP, Minghim R. 2015. InteractiVenn: a web-based tool for the analysis of sets through Venn diagrams. BMC Bioinformatics 16:169.

59. Ge SX, Jung D, Yao R. 2020. ShinyGO: a graphical gene-set enrichment tool for animals and plants. Bioinformatics 36:2628–2629.

